# Emergence and molecular basis of azithromycin resistance in typhoidal Salmonella in Dhaka, Bangladesh

**DOI:** 10.1101/594531

**Authors:** Yogesh Hooda, Senjuti Saha, Mohammad S. I. Sajib, Hafizur Rahman, Stephen P. Luby, Joseph Bondy-Denomy, Mathuram Santosham, Jason R. Andrews, Samir K. Saha

## Abstract

With rising fluoroquinolone and ceftriaxone-resistant *Salmonella* Typhi, azithromycin, a macrolide, has become the last oral drug available against typhoid. Between 2009-2016, we isolated 1,082 *Salmonella* Typhi and Paratyphi A strains in Bangladesh, 13 (12 Typhi and 1 Paratyphi A) of which were azithromycin-resistant. When compared to 462 previously sequenced Typhi strains, the genomes of the 12 azithromycin-resistant Typhi strains (4.3.1 sub-clade, H58) harbored an exclusive non-synonymous single-point mutation R717Q in AcrB, an RND-efflux pump. Expression of AcrB-R717Q in *E. coli* and Typhi strains increased its minimum inhibitory concentration (MIC) for azithromycin by 11- and 3-fold respectively. The azithromycin-resistant Paratyphi A strain also contained a mutation at R717 (R717L), whose introduction in *E. coli* and Paratyphi A strains increased MIC by 7- and 3-fold respectively, confirming the role of R717 mutations in conferring azithromycin resistance. With increasing azithromycin use, strains with R717 mutations may spread leading to treatment failures, making antibiotic stewardship and vaccine introduction imperative.

## Introduction

Typhoid and paratyphoid, collectively known as enteric fever, are among the most common bacterial causes of morbidity worldwide, with the greatest burden in low- and middle-income countries (GBD 2017 Typhoid and Paratyphoid Collaborators, 2019). *Salmonella enterica* subspecies *enterica* serovars Typhi (*Salmonella* Typhi) and Paratyphi (A, B and C), etiologies of enteric fever, cause an estimated 14 million illnesses and 136,000 deaths annually.

In the pre-antibiotic era, enteric fever mortality rates exceeded 20% in many areas, but ampicillin, chloramphenicol and co-trimoxazole were instrumental in reducing the rates to <1%. Resistance to all three antibiotics (referred to as multidrug resistance, MDR) emerged in late 1980’s (Mirza et al., 1996), predominantly due to the rise and subsequent continental migration of H58 haplotype (now referred to as 4.3.1), which contained the resistance genes either on IncH1 plasmids or integrated within the chromosome (Holt et al., 2011; Wong et al., 2015, 2016). Fluoroquinolones soon became the most-commonly prescribed antibiotic (White et al., 1996), but since the 2000’s there have been increasing reports of decreased fluoroquinolone susceptibility due to the acquisition of chromosomal mutations in the DNA gyrase and topoisomerase IV genes (Roumagnac et al., 2006; Chau et al., 2007, Dimitrov et al., 2007; Pham Thanh et al., 2016). In Bangladesh, >99% of all Typhi and Paratyphi strains exhibit decreased susceptibility to ciprofloxacin (Saha et al., 2018b). In 2011, WHO recommended ceftriaxone or azithromycin for treating *Salmonella* Typhi non-susceptible to fluroquinolones (Balasegaram et al., 2012).

There have been sporadic reports of ceftriaxone-resistant *Salmonella* Typhi strains, (Saha et al., 1999; Djeghout et al., 2018), but in 2016, an outbreak of extensively drug-resistant (XDR) *Salmonella* Typhi, resistant to chloramphenicol, ampicillin, cotrimoxazole, fluoroquinolones, and third-generation cephalosporins was recognized in Pakistan and to date >1000 cases have been confirmed (Andrews et al., 2018). Cephalosporin resistance of the XDR strains was caused by the acquisition of a broad-spectrum beta-lactamase (bla-CTX-M-15) on an IncY plasmid found in other enteric species. Typhoid patients in Pakistan are primarily being treated with the last available oral option, the macrolide azithromycin, resistance to which is rare (Klemm et al., 2018, Parry et al., 2015). This increasing use of azithromycin places selective pressure for the emergence and spread of azithromycin-resistant isolates, raising concerns of untreatable infections and increased mortality rates. Little is known about azithromycin resistance in typhoidal *Salmonella*; while there are some sporadic reports on azithromycin treatment failures (Molloy et al., 2010; Sjölund-Karlsson et al., 2011; Wong et al., 2015; Patel et al., 2017), there are no data on the molecular mechanism of resistance.

In Bangladesh, *Salmonella* Typhi and Paratyphi A are the most common causes of bloodstream infections in children >2 months of age and comprise of two-third of blood-culture positive isolates in microbiology laboratories (Saha et al., 2017). Leveraging our surveillance system in place for enteric fever, here we describe the emergence of azithromycin resistance among typhoidal *Salmonella* in Bangladesh and identify the molecular basis behind this resistance.

## Result and Discussion

### Emergence of azithromycin-resistant *Salmonella* Typhi and Paratyphi A

Between 2009 and 2016, through our enteric fever surveillance (Saha et al., 2017) in the inpatient departments of the two largest pediatric hospitals of Bangladesh, we isolated 939 *Salmonella* Typhi and 143 Paratyphi A strains. Twelve of the Typhi and one of the Paratyphi A strains were resistant to azithromycin, with disc diameters of ≤12 mm, and minimum inhibitory concentration (MIC) of ≥32 µg/ml (Parry et al., 2015). All 12 azithromycin-resistant *Salmonella* Typhi strains were also MDR and were increasingly isolated since 2013 (Fig 1A), marking gradual emergence of azithromycin-resistant *Salmonella* Typhi in Bangladesh. All patients lived in Dhaka city, known to be endemic for typhoid (Figure S1).

**Figure 1:**
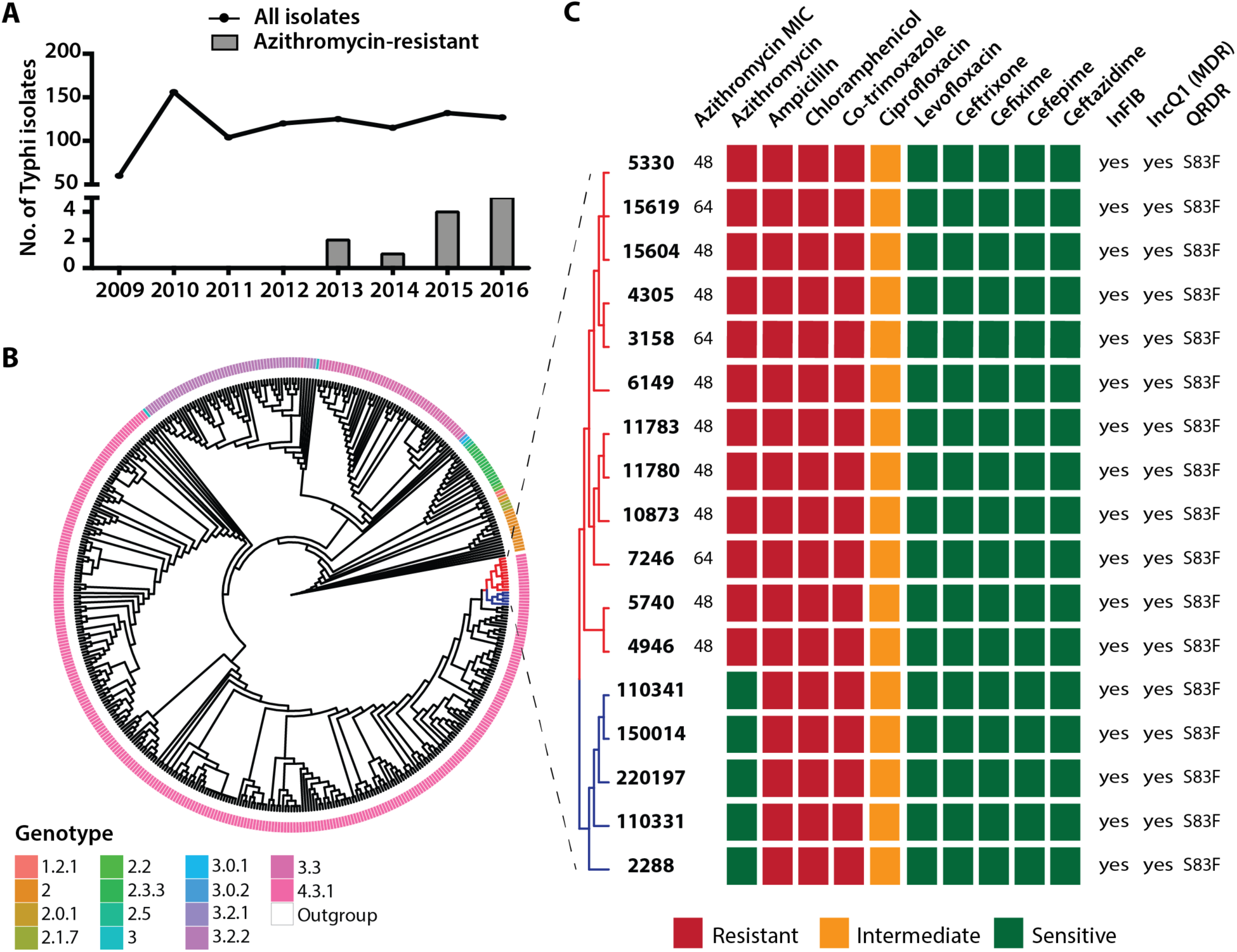
Emergence of azithromycin-resistant strains of *Salmonella* Typhi in Bangladesh and their genomic analysis. **A)** Temporal distribution of 939 *Salmonella* Typhi isolates included in the study. The number of isolates is shown as the line plot from 2009-2016. The numbers of azithromycin-resistant strains isolated each year is shown in the bar plot. Azithromycin-resistant strains were first isolated in 2013. **B)** Whole-genome SNP tree of 474 *Salmonella* Typhi strains isolated in Bangladesh previously (Tanmoy et al., 2018). The tree highlights the different genotypes that are found in Bangladesh including the most prevalent genotype 4.3.1 (H58 haplotype). The 12 azithromycin-resistant strains (colored in red) clustered together within the genotype 4.3.1. *Salmonella* typhimurium strain LT2 was used as an outgroup. **C)** Predicted and experimentally determined antimicrobial susceptibility pattern of azithromycin-resistant *Salmonella* Typhi strains and the most-closely related five azithromycin-sensitive strains. The antimicrobial susceptibility was experimentally determined through disc diffusion assay against a panel of 10 antibiotics. The predicted transmissible elements and antimicrobial resistance markers are also shown.

### Azithromycin resistant *Salmonella* Typhi harbors a mutation in the AcrB efflux pump

We sequenced the 12 azithromycin-resistant Typhi strains and found that all azithromycin-resistant strains belonged to genotype 4.3.1 (H58), the most common genotype found in South Asia (Tanmoy et al., 2018; Wong et al., 2015). In a whole-genome single nucleotide polymorphism (SNP) tree, the 12 strains clustered together indicating that they are genetically similar to one another and potentially arose for a single common parental strain (Figure 1B). To identify the genetic basis of azithromycin resistance, we used three bioinformatic tools: SRST2 (Inouye et al., 2014), Resfinder (Zankari et al., 2012) and CARD (Jia et al., 2017) and to evaluate the results obtained from these tools, we tested antimicrobial susceptibility against a panel of nine other antibiotics (Figure 1C). While the tools successfully predicted the observed susceptibility patterns for the nine antibiotics, no known azithromycin resistance mechanism was identified (Figure 1C). Using PlasmidFinder (Carattoli et al., 2014) we identified two plasmids found in these *Salmonella* Typhi strains: (i) IncQ1 (12/12 strains), containing genes for ampicillin, co-trimoxazole and chloramphenicol resistance, and (ii) InFIB (12/12 strains), a plasmid commonly found in *Salmonella* Typhi strains (Kidgell et al., 2002; Park et al., 2018). Both these plasmids were also present in closely related azithromycin-sensitive strains. The lack of known azithromycin-resistance genes indicated a novel mechanism of azithromycin resistance in these strains.

We compared the sequences of these 12 azithromycin-resistant strains to that of 462 Typhi strains that we had previously sequenced and genetically characterized (Tanmoy et al., 2018). In the WGS SNP tree, we identified four unique SNPs, present only in the 12 azithromycin-resistant strains, three of which were non-synonymous: STY2741 (codes for purN, a glycinamidine ribonucleotide transformyltransferase), STY1399 (codes for a hypothetical protein) and STY0519 (codes for AcrB, an inner membrane permease) (Figure 2A, Figure S2). For the first two candidates, there is no evidence of their involvement in mediating antimicrobial resistance in the literature. However, the third gene, *acrB* is part of a trans-envelope resistance-nodulation-division (RND) efflux pump that has been previously reported to transport macrolides including azithromycin across the bacterial cell envelope, making it the most promising candidate (Nakashima et al., 2011). Mutations affecting expression of AcrB have been implicated in macrolide resistance in *Neisseria gonorrhoeae* (Wadsworth et al., 2018). Furthermore, laboratory mutagenesis studies in *Escherichia coli* have shown that mutations in *acrB* can lead to higher macrolide efflux thereby contributing to resistance (Ababou and Koronakis, 2016). The SNP observed in the 12 azithromycin-resistant *Salmonella* Typhi strains changed the arginine residue (R) at position 717 to a glutamine (Q) (Figure 2B). R717 is a conserved residue on the periplasmic cleft that acts as the entry portal for most drugs in AcrB (Figure 2C). In a previous mutagenesis study, substitution of the arginine residue with an alanine (R717A) was found to partially increase efflux of the macrolide erythromycin in *E. coli* (Yu et al., 2005).

**Figure 2:**
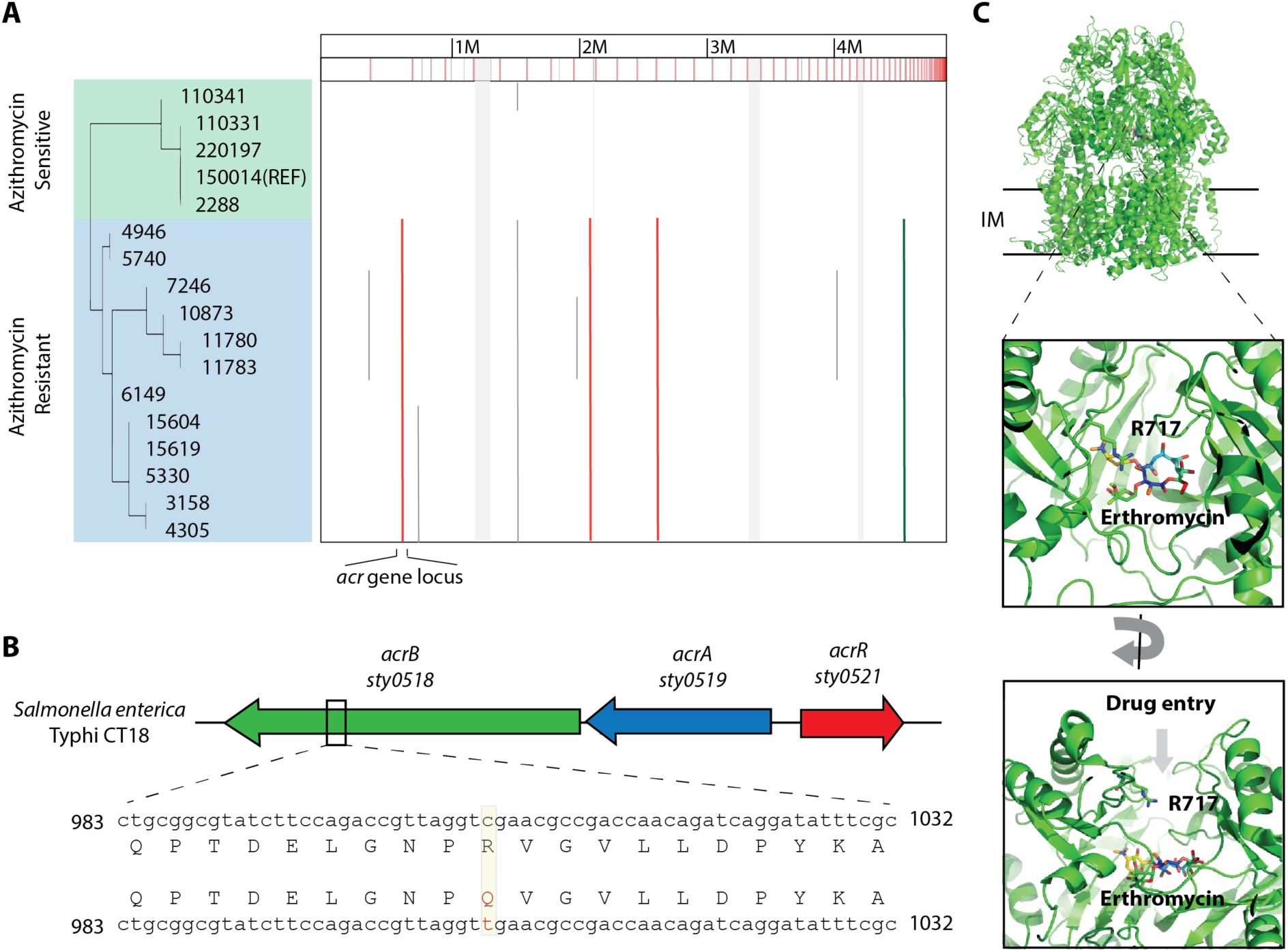
Identification of R717Q mutation on AcrB efflux pump as a cause of azithromycin resistance in *Salmonella* Typhi. **A)** Whole genome sequence alignment of 12 azithromycin-resistant and 5 genetically related azithromycin-sensitive *Salmonella* Typhi strains. Whole genome SNP detection and alignment was done using ParSNP and results were visualized in GinGR (Treangen et al., 2014). The single nucleotide polymorphisms (SNPs) unique to the resistant strains are highlighted with vertical lines. Four SNPs were identified: 3 non-synonymous (shown as a red line) and 1 synonymous SNPs (shown as a brown line) that are exclusive to the azithromycin resistant strains. **B)** The *acr* gene cluster in *Salmonella* Typhi reference strain CT18. One of the SNPs found exclusively in azithromycin-resistant strains was mapped to the gene cluster composed of: *acrA* (STY0519) and *acrB* (STY0518), that encodes a periplasmic and inner membrane protein component of the RND-efflux pumps respectively, and *acrR* (STY0521), a transcriptional regulator of AcrA/B protein synthesis. The SNP was present on the *acrB* gene and resulted in the change of an arginine (R) at position 717 to a glutamine (Q) residue on the encoded AcrB protein. **C)** R717Q mutation is present at the periplasmic cleft of the proximal binding pocket on AcrB. Structure of *E. coli* AcrB (PDB ID: 3AOC) is shown in green with the macrolide erythromycin bound in the proximal drug binding pocket. AcrB is present in the inner membrane of the bacterial cells and drug molecules, including macrolides, enter the AcrB pump through a periplasmic opening that leads to a proximal binding pocket. The drug molecules are shuttled outside the cells through the proximal binding pocket with the help of the proton motive force. R717 lines the entry the periplasmic opening.

### R717 mutations in AcrB confer azithromycin resistance

We cloned *acrB* from azithromycin susceptible and resistant *Salmonella* Typhi strains into an *E. coli* plasmid and introduced them into an *E. coli* strain that lacks the endogenous *acrB* (*E. coli ΔacrB*). Compared to *E. coli* strains containing empty plasmid or wild type *acrB*, the strain expressing AcrB-R717Q showed a smaller zone of disc clearance for both azithromycin (26.3 mm vs 16.7 mm, *p* = 0.0013) and erythromycin discs (22.3 mm vs 11.7 mm, *p value* = 0.0009) and exhibited a 11-fold increase in azithromycin MIC (0.22 µg/ml vs 2.7 µg/ml, *p* = 0.0002) (Figure 3A, B). Certain AcrB mutations have previously been shown to effect transport of other antibiotics such as ciprofloxacin (Blair et al., 2015), but the R717Q mutation did not change the susceptibility patterns for any other nine antibiotics we tested (Figure S3). For further confirmation of the effects of this mutation in *Salmonella* Typhi, we introduced the plasmids in an azithromycin-sensitive Typhi strain and observed a 3-fold increase in MIC (5 µg/ml vs 13 µg/ml, *p* < 0.0001) in the presence of AcrB-R717Q (Figure 3C). The difference here is lower compared to that seen in *E. coli ΔacrB* plausibly because the Typhi strain contains endogenous wild-type AcrB competing against the exogenous AcrB-R717Q that we artificially introduced. Taken together, these results confirm that AcrB-R717Q leads to increased macrolide resistance.

**Figure 3:**
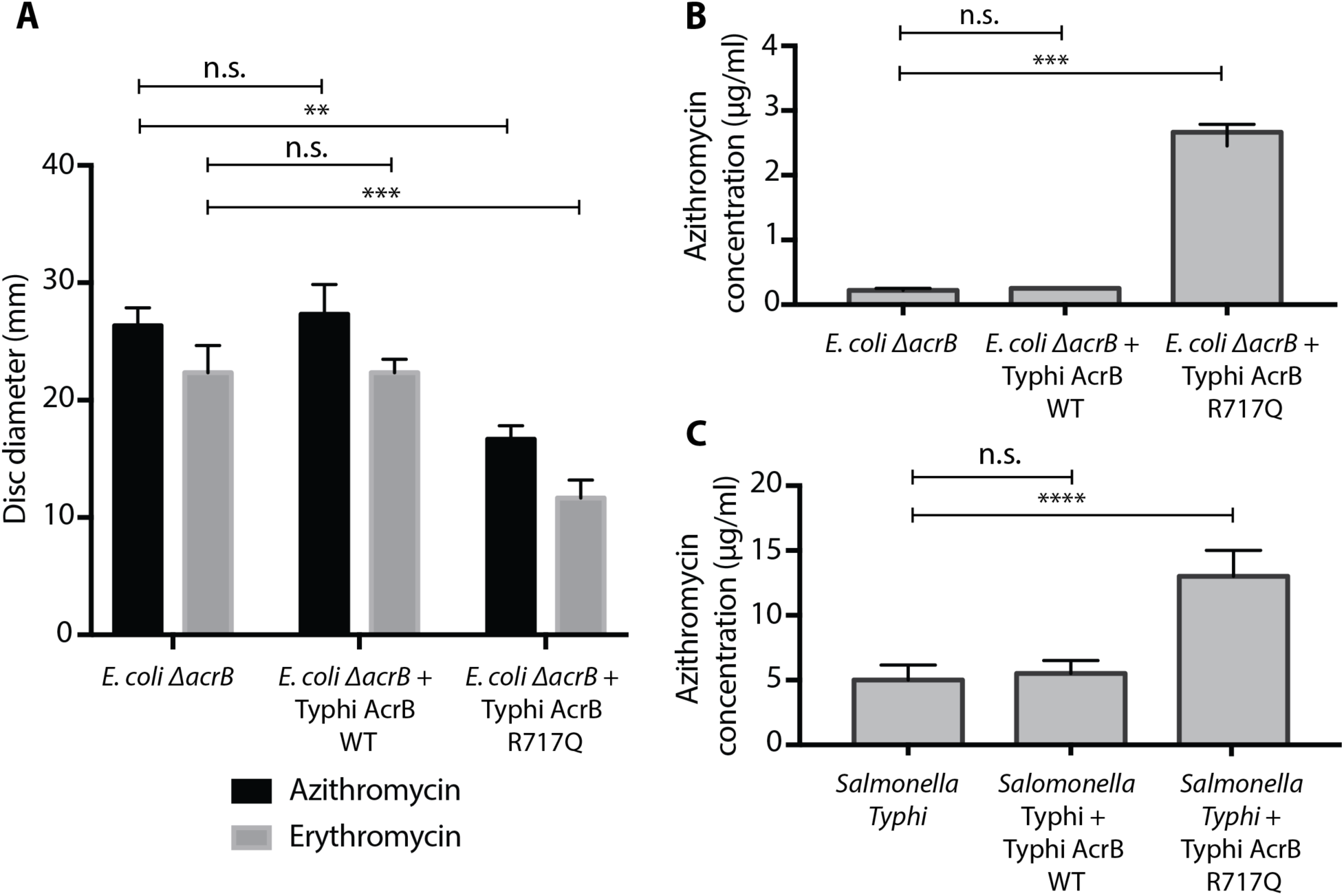
AcrB R717Q increases efflux of macrolides in *E*. *coli* and *Salmonella* Typhi strains. Quantification of results obtained from three biological replicates for **(A)** disc diffusion and **(B)** E-strip assays are shown for *E. coli* BW25113 *ΔacrB* transformed with different plasmids. **(C)** Quantification of results obtained from at least three biological replicates from azithromycin E-strip assay in Typhi strain 4119. One way-ANOVA with multiple comparisons was used to test statistical significance. ns: p>0.05; **: p ≤ 0.01; ***: p ≤ 0.001, ****: p ≤ 0.0001

We conducted an extensive BLAST search to identify other typhoidal *Salmonella* strains with mutations in AcrB the NCBI database and found only one *Salmonella* Typhi strain isolated in Oceania in 2008 that contained the same R717Q mutation, however no AMR data were available for this strain (Wong et al., 2015). Interestingly, whole genome sequencing of the one azithromycin-resistant *Salmonella* Paratyphi A strain identified during our surveillance showed that this strain also contained a mutation in *acrB* which changed R717 to a leucine (L) (Figure 4A). This mutation was absent in the genomes of the Paratyphi A strains in the NCBI database. To determine to effect of R717L mutation, we expressed Paratyphi wild-type AcrB and AcrB-R717L in *E. coli ΔacrB* (Figure 4B, C). As seen for Typhi AcrB R717Q, Paratyphi AcrB-R717L leads to a smaller disc clearance for azithromycin (26.3 mm vs 16.3 mm, *p* = 0.0001) and erythromycin (22.4 mm vs 11.4 mm, *p* = 0.0001) and 10-fold higher azithromycin MIC (0.22 µg/ml vs 2.5 µg/ml, *p* = 0.003). When these plasmids were introduced in a sensitive Paratyphi A strain, we observed 4-fold change in MIC (7 µg/ml vs 28 µg/ml, *p* = 0.0001) in the presence of the R7171L mutation, confirming that mutations in R717 lead to macrolide resistance.

**Figure 4:**
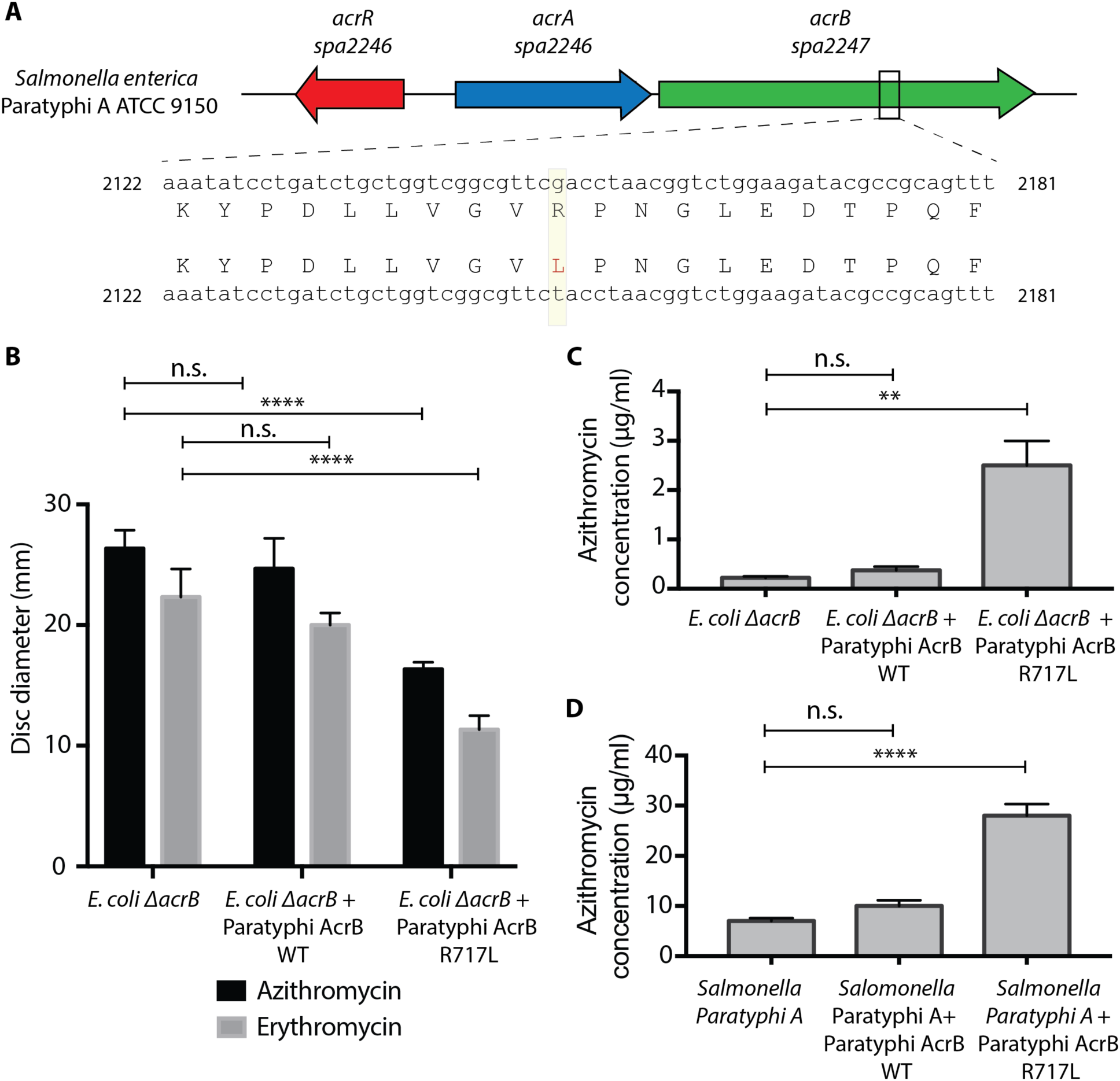
Identification of R717L mutation in AcrB protein in *Salmonella* Paratyphi A strains. **(A)** Sequence alignment of *acrB* gene from the azithromycin-resistant *Salmonella* Paratyphi A strain to the reference strain ATCC 9150. A SNP was identified that changed the R717 to a leucine (L) residue. Quantification of results obtained from three biological replicates for **(B)** disc diffusion and **(C)** E-strip assays in *E. coli* are shown. **(D)** Quantification of results obtained from at least three biological replicates from azithromycin E-strip assay in Paratyphi strain 4071 is shown. One way-ANOVA with multiple comparisons was used to test statistical significance. ns: p>0.05; **: p ≤ 0.01; ****: p ≤ 0.0001.

Rising antimicrobial resistance threatens the progress made so far in management of enteric fever. The rate of azithromycin resistance of typhoidal *Salmonella* in Bangladesh is low and the genetic basis is a chromosomal SNP. However, in light of the outbreak in Pakistan, increasing azithromycin use can place selective pressure on strains such as the ones with R717 mutations to spread. Although no azithromycin resistant XDR isolate has been reported to date, the increasing use of azithromycin and the clear historical record of widespread dissemination of resistance to all previously widely used antimicrobials by *Salmonella* Typhi and Paratyphi, suggest we will soon face strains resistant to all oral antibiotics. An azithromycin-resistant XDR strain would shift enteric fever treatment from outpatient departments, where patients are currently treated with oral azithromycin, to inpatient departments to be treated with injectable antibiotics like carbapenems, thereby weighing down already struggling health systems in endemic regions (Andrews et al., 2018; Saha et al., 2018a). Moreover, with the dearth of novel antimicrobials in the horizon, we risk losing our primary defense against widespread mortality from enteric fever. In addition to continued surveillance and antimicrobial stewardship, it is imperative to roll-out the recent WHO prequalified typhoid conjugate vaccine in endemic areas and decrease the overall burden of typhoid.

## Materials and Methods

### Study site and population

In this study, we report data from enteric fever surveillance conducted in the inpatient departments of the two largest pediatric hospitals of Bangladesh, Dhaka Shishu (Children) Hospital, DSH, and Shishu Shasthya (Child Health) Foundation Hospital, SSFH. These are sentinel sites of the World Health Organization supported Invasive Bacterial Vaccine Preventable Diseases surveillance platform in Bangladesh.

### Patient enrollment, etiology detection and antibiogram

Blood culture was performed at the discretion of the treating physicians. We enrolled patients with positive blood cultures for *Salmonella* Typhi or Paratyphi A. Blood cultures were performed using standard methods (Saha et al., 2017). We aseptically obtained 2–3 milliliters of blood, which was inoculated into trypticase soy broth supplemented with sodium polyanethole sulphonate (0.25%) and isovitalex (1%). Incubated blood culture bottles were sub-cultured on the second, third, and fifth days of incubation. Identification of *Salmonella* Typhi/Paratyphi isolates was confirmed using standard biochemical tests and agglutination with *Salmonella* species and serovar-specific antisera (Ramel, Thermo Fisher Scientific). Laboratory methods for blood culture and organism identification were consistent over the reporting period.

We used disc diffusion methods for determining antibiotic susceptibility patterns for azithromycin, ampicillin, co-trimoxazole, chloramphenicol, ciprofloxacin, levofloxacin, ceftriaxone, cefepime, cefixime and ceftazidime (Oxoid, Thermo Scientific, MA, USA). Azithromycin e-strips (bioMérieux, France) were used to determine the minimum inhibitory concentration (MIC) and confirm azithromycin resistance for strains that exhibited zone of clearance ≤12 mm in the presence of azithromycin discs. All results were interpreted according to the latest Clinical and Laboratory Standards Institute guidelines 2018.

### DNA extraction and whole genome sequencing

We conducted whole genome sequencing on all identified azithromycin-resistant strains (12 *Salmonella* Typhi and 1 Paratyphi A). Isolates were grown in MacConkey agar (Oxoid, UK) overnight and DNA was extracted from a suspension of the overnight culture using the QIAamp DNA minikit (Qiagen, Hilden, Germany). Whole genome sequencing was performed on the Illumina HiSeq 4000 platform to generate 150 bp paired-end reads (Novogene Co. Ltd., Beijing, China). We used SPAdes (Bankevich et al., 2012) to assemble the short paired-end reads into contigs for downstream analyses. All the sequences have been submitted to EnteroBase and NCBI (BioProject ID: PRJNA528114).

### Bioinformatics analysis

For comparative genomic analysis, we compared the 12 azithromycin-resistant *Salmonella* Typhi strains with 462 strains that were previously isolated and genetically characterized by our group in Bangladesh (Tanmoy et al., 2018). Using the ParSNP tool (Treangen et al., 2014), we constructed whole-genome SNP tree using all 475 genomes and determined the genotypes using srst2 package (Inouye et al., 2014). Ggtree was used to make the phylogenetic tree and overlay the genotype data (Yu et al., 2017). Srst2, ResFinder (Zankari et al., 2012) and CARD (Jia et al., 2017) were used to predict antimicrobial resistance markers, and PlasmidFinder (Carattoli et al., 2014) to identify the putative plasmids present in these strains. Finally, we compared the resistant strains to all sensitive *Salmonella* Typhi strains manually to find SNPs exclusive to the resistant strains (comparison to the most closely related 5 genomes are shown in Figure 1C) using the GinGR tool from the Harvest suite (Treangen et al., 2014). To predict the function of the SNPs on protein function, we examined the protein sequence and conducted structural analyses (Figure 2).

### Macrolide susceptibility test in *E*. *coli* and *Salmonella* Typhi

We amplified the *acrB* genes from azithromycin-resistant *Salmonella* Typhi strain 5003 and Paratyphi A strain 3144 and azithromycin-sensitive Typhi strain 4119 and Paratyphi A strain 4071 for downstream cloning. The genes were inserted into the multiple cloning site of pHERD30T using Gibson assembly (Gibson et al., 2009). We verified the sequences of all inserted genes through Sanger sequencing. The plasmids were chemically transformed into CaCl_2_-competent *E. coli* BW25112 *ΔacrB* strains, and electroporated into Typhi strain 4119 and Paratyphi A strain 4071. *E. coli* strains with plasmids (with or without an insert) were tested for susceptibility patterns for erythromycin, azithromycin, ampicillin, co-trimoxazole, chloramphenicol, ciprofloxacin, levofloxacin, ceftriaxone, cefepime, cefixime and ceftazidime using the disc diffusion method, and MIC was determined using azithromycin E-strips. Typhi and Paratyphi strains with plasmids (with or without an insert) were tested for susceptibility patterns for azithromycin using E-strips.

## Acknowledgements

We are thankful to Dr. Shweta Karambelkar, Dr. Balint Csorgo, and Beatriz A. Osuna of University of California, San Francisco, and Sunita Patil of Stanford University for technical assistance with laboratory work and to Arif M. Tanmoy of the Child Health Research Foundation for bioinformation guidance.

## Supplementary Figures

**Figure S1:**
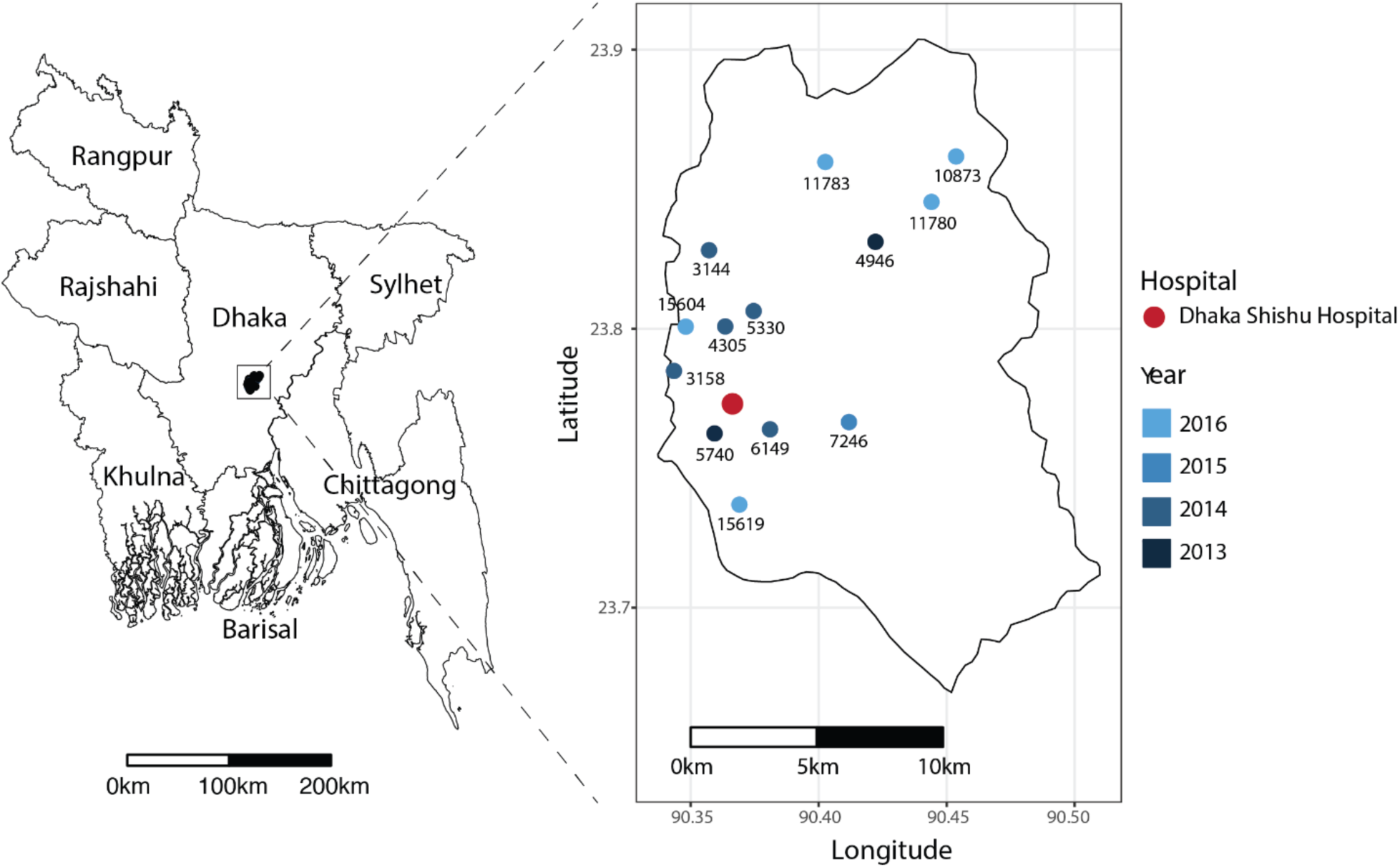
Spatiotemporal distribution of azithromycin-resistant *Salmonella* Typhi and Paratyphi A strains. The 13 azithromycin-resistant typhoidal *Salmonella* strains were isolated from Dhaka Shishu Hospital (shown in red). All the patients lived within the Dhaka municipal area.

**Figure S2:**
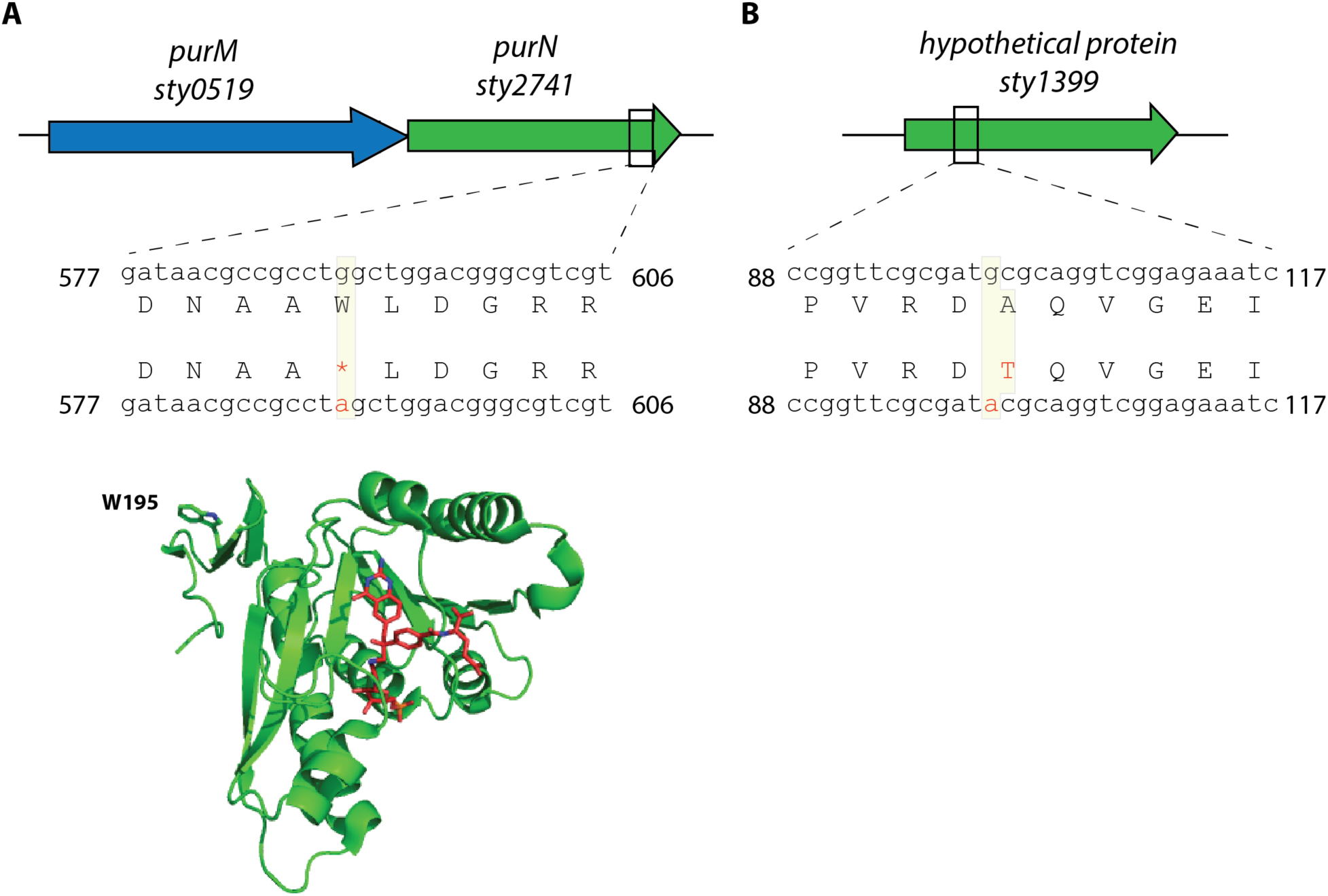
Genetic and structural analysis of 2 other non-synonymous SNPs. **A)** SNP on *sty2741* gene (also known as *purN*) that encodes a glycinamidine ribonucleotide transformyltransferase (GAR-Tfase) enzyme. The SNP leads to change in W195 to a stop codon, leading to premature termination of the protein sequence. The structure of *E. coli* GAT-Tfase (green, PDB ID: 1C3E) in complex with an inhibitor (shown in red) highlighting the active site is shown. The W195 is present close to the C-terminus and premature termination at this position is predicted to not affect protein function. **C)** SNP on *sty1399* that encodes a hypothetical protein that is proposed to contain a B3/B4 tRNA-binding domain. The function of this protein is not known and the SNP results in conversion of an alanine residue at position 34 to a threonine residue. None of these two genes have been previously implicated in macrolide resistance.

**Figure S3 – figure supplement 3:**
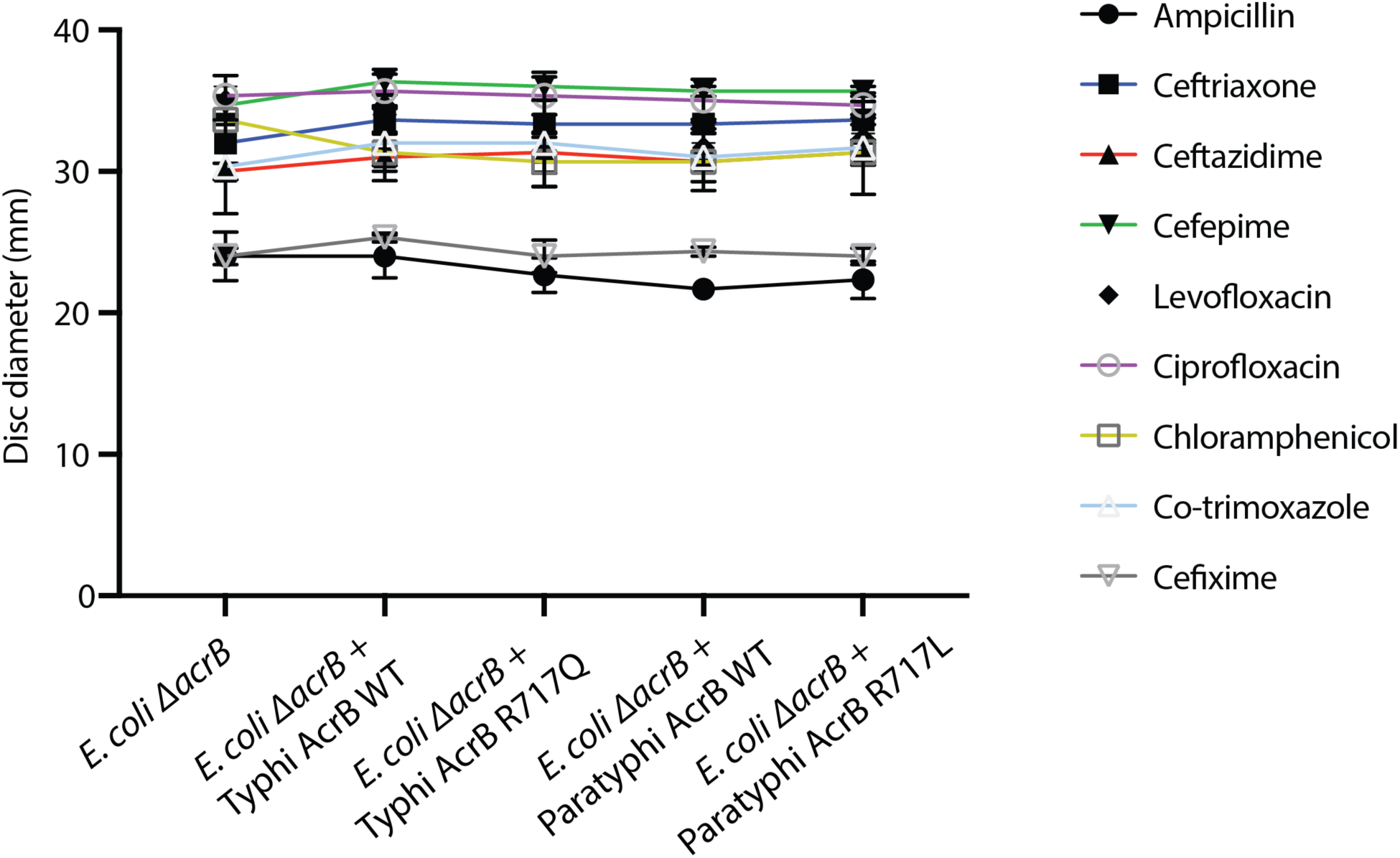
AcrB mutations do not affect efflux of other families of antibiotics. Susceptibility of *E. coli* strains with empty, *Salmonella* Typhi AcrB WT/R717Q and Paratyphi A AcrB WT/R717L were tested against a panel of 9 different antibiotic including 5 beta-lactams, 2 fluoroquinolones, 1 phenicol and 1 diaminopyrimidine /sulphonamide. The data are shown as mean and standard error from 3 different biological replicates.

## References

Ababou A, Koronakis V. 2016. Structures of Gate Loop Variants of the AcrB Drug Efflux Pump Bound by Erythromycin Substrate. PLOS ONE 11:e0159154. doi:10.1371/journal.pone.0159154

Andrews JR, Qamar FN, Charles RS, Ryan ET. 2018. Extensively Drug-Resistant Typhoid — Are Conjugate Vaccines Arriving Just in Time? NEJM 379:1493–1495.

Balasegaram S, Potter AL, Grynszpan D, Barlow S, Behrens RH, Lighton L, Booth L, Inamdar L, Neal K, Nye K, Lawrence J, Jones J, Gray I, Tolley D, Lane C, Adak B, Cummins A, Addiman S. 2012. Guidelines for the public health management of typhoid and paratyphoid in England: Practice guidelines from the National Typhoid and Paratyphoid Reference Group. Journal of Infection 65:197–213. doi:10.1016/j.jinf.2012.05.005

Bankevich A, Nurk S, Antipov D, Gurevich AA, Dvorkin M, Kulikov AS, Lesin VM, Nikolenko SI, Pham S, Prjibelski AD, Pyshkin AV, Sirotkin AV, Vyahhi N, Tesler G, Alekseyev MA, Pevzner PA. 2012. SPAdes: a new genome assembly algorithm and its applications to single-cell sequencing. J Comput Biol 19:455–477. doi:10.1089/cmb.2012.0021

Blair JMA, Bavro VN, Ricci V, Modi N, Cacciotto P, Kleinekath?fer U, Ruggerone P, Vargiu AV, Baylay AJ, Smith HE, Brandon Y, Galloway D, Piddock LJV. 2015. AcrB drug-binding pocket substitution confers clinically relevant resistance and altered substrate specificity. PNAS 112:3511–3516. doi:10.1073/pnas.1419939112

Carattoli A, Zankari E, García-Fernández A, Larsen MV, Lund O, Villa L, Aarestrup FM, Hasman H. 2014. In Silico Detection and Typing of Plasmids using PlasmidFinder and Plasmid Multilocus Sequence Typing. Antimicrobial Agents and Chemotherapy 58:3895–3903. doi:10.1128/AAC.02412-14

Chau TT, Campbell JI, Galindo CM, Van Minh Hoang N, Diep TS, Nga TTT, Van Vinh Chau N, Tuan PQ, Page AL, Ochiai RL, Schultsz C, Wain J, Bhutta ZA, Parry CM, Bhattacharya SK, Dutta S, Agtini M, Dong B, Honghui Y, Anh DD, Canh DG, Naheed A, Albert MJ, Phetsouvanh R, Newton PN, Basnyat B, Arjyal A, La TTP, Rang NN, Phuong LT, Van Be Bay P, von Seidlein L, Dougan G, Clemens JD, Vinh H, Hien TT, Chinh NT, Acosta CJ, Farrar J, Dolecek C. 2007. Antimicrobial Drug Resistance of Salmonella enterica Serovar Typhi in Asia and Molecular Mechanism of Reduced Susceptibility to the Fluoroquinolones. Antimicrob Agents Chemother 51:4315–4323. doi:10.1128/AAC.00294-07

Dimitrov T, Udo EE, Albaksami O, Kilani AA, Shehab E-DMR. 2007. Ciprofloxacin treatment failure in a case of typhoid fever caused by Salmonella enterica serotype Paratyphi A with reduced susceptibility to ciprofloxacin. Journal of Medical Microbiology 56:277–279. doi:10.1099/jmm.0.46773-0

Djeghout B, Saha S, Sajib MSI, Tanmoy AM, Islam M, Kay GL, Langridge GC, Endtz HP, Wain J, Saha SK. 2018. Ceftriaxone-resistant Salmonella Typhi carries an IncI1-ST31 plasmid encoding CTX-M-15. Journal of Medical Microbiology 67:620–627. doi:10.1099/jmm.0.000727

GBD 2017 Typhoid and Paratyphoid Collaborators. 2019. The global burden of typhoid and paratyphoid fevers: a systematic analysis for the Global Burden of Disease Study 2017. Lancet Infect Dis. doi:10.1016/S1473-3099(18)30685-6

Gibson DG, Young L, Chuang R-Y, Venter JC, Hutchison Iii CA, Smith HO. 2009. Enzymatic assembly of DNA molecules up to several hundred kilobases. Nature Methods 6:343–345. doi:10.1038/nmeth.1318

Holt KE, Phan MD, Baker S, Duy PT, Nga TVT, Nair S, Turner AK, Walsh C, Fanning S, Farrell-Ward S, Dutta S, Kariuki S, Weill F-X, Parkhill J, Dougan G, Wain J. 2011. Emergence of a Globally Dominant IncHI1 Plasmid Type Associated with Multiple Drug Resistant Typhoid. PLoS Negl Trop Dis 5. doi:10.1371/journal.pntd.0001245

Inouye M, Dashnow H, Raven L-A, Schultz MB, Pope BJ, Tomita T, Zobel J, Holt KE. 2014. SRST2: Rapid genomic surveillance for public health and hospital microbiology labs. Genome Medicine 6:90. doi:10.1186/s13073-014-0090-6

Jia B, Raphenya AR, Alcock B, Waglechner N, Guo P, Tsang KK, Lago BA, Dave BM, Pereira S, Sharma AN, Doshi S, Courtot M, Lo R, Williams LE, Frye JG, Elsayegh T, Sardar D, Westman EL, Pawlowski AC, Johnson TA, Brinkman FSL, Wright GD, McArthur AG. 2017. CARD 2017: expansion and model-centric curation of the comprehensive antibiotic resistance database. Nucleic Acids Res 45:D566–D573. doi:10.1093/nar/gkw1004

Kidgell C, Pickard D, Wain J, James K, Diem Nga LT, Diep TS, Levine MM, O’Gaora P, Prentice MB, Parkhill J, Day N, Farrar J, Dougan G. 2002. Characterisation and distribution of a cryptic Salmonella typhi plasmid pHCM2. Plasmid 47:159–171. doi:10.1016/S0147-619X(02)00013-6

Klemm EJ, Shakoor S, Page AJ, Qamar FN, Judge K, Saeed DK, Wong VK, Dallman TJ, Nair S, Baker S, Shaheen G, Qureshi S, Yousafzai MT, Saleem MK, Hasan Z, Dougan G, Hasan R. 2018. Emergence of an Extensively Drug-Resistant Salmonella enterica Serovar Typhi Clone Harboring a Promiscuous Plasmid Encoding Resistance to Fluoroquinolones and Third-Generation Cephalosporins. mBio 9:e00105–18. doi:10.1128/mBio.00105-18

Mirza SH, Beechmg NJ, Hart CA. 1996. Multi-drug resistant typhoid: a global problem. Journal of Medical Microbiology 44:317–319. doi:10.1099/00222615-44-5-317

Molloy A, Nair S, Cooke FJ, Wain J, Farrington M, Lehner PJ, Torok ME. 2010. First Report of Salmonella enterica Serotype Paratyphi A Azithromycin Resistance Leading to Treatment Failure. Journal of Clinical Microbiology 48:4655–4657. doi:10.1128/JCM.00648-10

Nakashima R, Sakurai K, Yamasaki S, Nishino K, Yamaguchi A. 2011. Structures of the multidrug exporter AcrB reveal a proximal multisite drug-binding pocket. Nature 480:565–569. doi:10.1038/nature10641

Park SE, Pham DT, Boinett C, Wong VK, Pak GD, Panzner U, Espinoza LMC, Kalckreuth V von, Im J, Schütt-Gerowitt H, Crump JA, Breiman RF, Adu-Sarkodie Y, Owusu-Dabo E, Rakotozandrindrainy R, Soura AB, Aseffa A, Gasmelseed N, Keddy KH, May J, Sow AG, Aaby P, Biggs HM, Hertz JT, Montgomery JM, Cosmas L, Olack B, Fields B, Sarpong N, Razafindrabe TJL, Raminosoa TM, Kabore LP, Sampo E, Teferi M, Yeshitela B, Tayeb MAE, Sooka A, Meyer CG, Krumkamp R, Dekker DM, Jaeger A, Poppert S, Tall A, Niang A, Bjerregaard-Andersen M, Løfberg SV, Seo HJ, Jeon HJ, Deerin JF, Park J, Konings F, Ali M, Clemens JD, Hughes P, Sendagala JN, Vudriko T, Downing R, Ikumapayi UN, Mackenzie GA, Obaro S, Argimon S, Aanensen DM, Page A, Keane JA, Duchene S, Dyson Z, Holt KE, Dougan G, Marks F, Baker S. 2018. The phylogeography and incidence of multi-drug resistant typhoid fever in sub-Saharan Africa. Nature Communications 9:5094. doi:10.1038/s41467-018-07370-z

Parry CM, Thieu NTV, Dolecek C, Karkey A, Gupta R, Turner P, Dance D, Maude RR, Ha V, Tran CN, Thi PL, Be BPV, Phi LTT, Ngoc RN, Ghose A, Dongol S, Campbell JI, Thanh DP, Thanh TH, Moore CE, Sona S, Gaind R, Deb M, Anh HV, Van SN, Tinh HT, Day NPJ, Dondorp A, Thwaites G, Faiz MA, Phetsouvanh R, Newton P, Basnyat B, Farrar JJ, Baker S. 2015. Clinically and Microbiologically Derived Azithromycin Susceptibility Breakpoints for Salmonella enterica Serovars Typhi and Paratyphi A. Antimicrobial Agents and Chemotherapy 59:2756–2764. doi:10.1128/AAC.04729-14

Patel SR, Bharti S, Pratap CB, Nath G. 2017. Drug Resistance Pattern in the Recent Isolates of Salmonella Typhi with Special Reference to Cephalosporins and Azithromycin in the Gangetic Plain. J Clin Diagn Res 11:DM01–DM03. doi:10.7860/JCDR/2017/23330.9973

Pham Thanh D, Karkey A, Dongol S, Ho Thi N, Thompson CN, Rabaa MA, Arjyal A, Holt KE, Wong V, Tran Vu Thieu N, Voong Vinh P, Ha Thanh T, Pradhan A, Shrestha SK, Gajurel D, Pickard D, Parry CM, Dougan G, Wolbers M, Dolecek C, Thwaites GE, Basnyat B, Baker S. 2016. A novel ciprofloxacin-resistant subclade of H58 Salmonella Typhi is associated with fluoroquinolone treatment failure. eLife 5. doi:10.7554/eLife.14003

Roumagnac P, Weill F-X, Dolecek C, Baker S, Brisse S, Chinh NT, Le TAH, Acosta CJ, Farrar J, Dougan G, Achtman M. 2006. Evolutionary History of Salmonella Typhi. Science 314:1301–1304. doi:10.1126/science.1134933

Saha S, Santosham M, Hussain M, Black RE, Saha SK. 2018a. Rotavirus Vaccine will Improve Child Survival by More than Just Preventing Diarrhea: Evidence from Bangladesh. The American Journal of Tropical Medicine and Hygiene 98:360–363. doi:10.4269/ajtmh.17-0586

Saha Senjuti, Islam M, Saha Shampa, Uddin MJ, Rahman H, Das RC, Hasan M, Amin MR, Hanif M, Shahidullah M, Hussain M, Saha SK. 2018b. Designing Comprehensive Public Health Surveillance for Enteric Fever in Endemic Countries: Importance of Including Different Healthcare Facilities. J Infect Dis 218:S227–S231. doi:10.1093/infdis/jiy191

Saha Senjuti, Islam M, Uddin MJ, Saha Shampa, Das RC, Baqui AH, Santosham M, Black RE, Luby SP, Saha SK. 2017. Integration of enteric fever surveillance into the WHO-coordinated Invasive Bacterial-Vaccine Preventable Diseases (IB-VPD) platform: A low cost approach to track an increasingly important disease. PLOS Neglected Tropical Diseases 11:e0005999. doi:10.1371/journal.pntd.0005999

Saha SK, Talukder SY, Islam M, Saha S. 1999. A highly ceftriaxone-resistant Salmonella typhi in Bangladesh. Pediatr Infect Dis J 18:387.

Sjölund-Karlsson M, Joyce K, Blickenstaff K, Ball T, Haro J, Medalla FM, Fedorka-Cray P, Zhao S, Crump JA, Whichard JM. 2011. Antimicrobial Susceptibility to Azithromycin among Salmonella enterica Isolates from the United States?. Antimicrob Agents Chemother 55:3985–3989. doi:10.1128/AAC.00590-11

Tanmoy AM, Westeel E, Bruyne KD, Goris J, Rajoharison A, Sajib MSI, Belkum A van, Saha SK, Komurian-Pradel F, Endtz HP. 2018. Salmonella enterica Serovar Typhi in Bangladesh: Exploration of Genomic Diversity and Antimicrobial Resistance. mBio 9:e02112–18. doi:10.1128/mBio.02112-18

Treangen TJ, Ondov BD, Koren S, Phillippy AM. 2014. The Harvest suite for rapid core-genome alignment and visualization of thousands of intraspecific microbial genomes. Genome Biology 15:524. doi:10.1186/s13059-014-0524-x

Wadsworth CB, Arnold BJ, Sater MRA, Grad YH. 2018. Azithromycin Resistance through Interspecific Acquisition of an Epistasis-Dependent Efflux Pump Component and Transcriptional Regulator in Neisseria gonorrhoeae. mBio 9:e01419–18. doi:10.1128/mBio.01419-18

White NJ, Dung NM, Vinh H, Bethell D, Hien IT. 1996. Fluoroquinolone antibiotics in children with multidrug resistant typhoid. The Lancet 348:547. doi:10.1016/S0140-6736(05)64703-4

Wong VK, Baker S, Connor TR, Pickard D, Page AJ, Dave J, Murphy N, Holliman R, Sefton A, Millar M, Dyson ZA, Dougan G, Holt KE, International Typhoid Consortium, Parkhill J, Feasey NA, Kingsley RA, Thomson NR, Keane JA, Weill F-X, Le Hello S, Hawkey J, Edwards DJ, Harris SR, Cain AK, Hadfield J, Hart PJ, Thieu NTV, Klemm EJ, Breiman RF, Watson CH, Edmunds WJ, Kariuki S, Gordon MA, Heyderman RS, Okoro C, Jacobs J, Lunguya O, Msefula C, Chabalgoity JA, Kama M, Jenkins K, Dutta S, Marks F, Campos J, Thompson C, Obaro S, MacLennan CA, Dolecek C, Keddy KH, Smith AM, Parry CM, Karkey A, Dongol S, Basnyat B, Arjyal A, Mulholland EK, Campbell JI, Dufour M, Bandaranayake D, Toleafoa TN, Singh SP, Hatta M, Newton PN, Dance D, Davong V, Onsare RS, Isaia L, Thwaites G, Wijedoru L, Crump JA, De Pinna E, Nair S, Nilles EJ, Thanh DP, Turner P, Soeng S, Valcanis M, Powling J, Dimovski K, Hogg G, Farrar J, Mather AE, Amos B. 2016. An extended genotyping framework for *Salmonella enterica* serovar Typhi, the cause of human typhoid. Nature Communications 7:12827. doi:10.1038/ncomms12827

Wong VK, Baker S, Pickard DJ, Parkhill J, Page AJ, Feasey NA, Kingsley RA, Thomson NR, Keane JA, Weill F-X, Edwards DJ, Hawkey J, Harris SR, Mather AE, Cain AK, Hadfield J, Hart PJ, Thieu NTV, Klemm EJ, Glinos DA, Breiman RF, Watson CH, Kariuki S, Gordon MA, Heyderman RS, Okoro C, Jacobs J, Lunguya O, Edmunds WJ, Msefula C, Chabalgoity JA, Kama M, Jenkins K, Dutta S, Marks F, Campos J, Thompson C, Obaro S, MacLennan CA, Dolecek C, Keddy KH, Smith AM, Parry CM, Karkey A, Mulholland EK, Campbell JI, Dongol S, Basnyat B, Dufour M, Bandaranayake D, Naseri TT, Singh SP, Hatta M, Newton P, Onsare RS, Isaia L, Dance D, Davong V, Thwaites G, Wijedoru L, Crump JA, De Pinna E, Nair S, Nilles EJ, Thanh DP, Turner P, Soeng S, Valcanis M, Powling J, Dimovski K, Hogg G, Farrar J, Holt KE, Dougan G. 2015. Phylogeographical analysis of the dominant multidrug-resistant H58 clade of Salmonella Typhi identifies inter-and intracontinental transmission events. Nat Genet 47:632–639. doi:10.1038/ng.3281

Yu EW, Aires JR, McDermott G, Nikaido H. 2005. A Periplasmic Drug-Binding Site of the AcrB Multidrug Efflux Pump: a Crystallographic and Site-Directed Mutagenesis Study. J Bacteriol 187:6804–6815. doi:10.1128/JB.187.19.6804-6815.2005

Yu G, Smith DK, Zhu H, Guan Y, Lam TT-Y. 2017. ggtree: an r package for visualization and annotation of phylogenetic trees with their covariates and other associated data. Methods in Ecology and Evolution 8:28–36. doi:10.1111/2041-210X.12628

Zankari E, Hasman H, Cosentino S, Vestergaard M, Rasmussen S, Lund O, Aarestrup FM, Larsen MV. 2012. Identification of acquired antimicrobial resistance genes. J Antimicrob Chemother 67:2640–2644. doi:10.1093/jac/dks261

